# Gene regulation of the avian malaria parasite *Plasmodium relictum*, during the different stages within the mosquito vector

**DOI:** 10.1101/2020.07.16.204198

**Authors:** V. Sekar, A. Rivero, R. Pigeault, S. Gandon, A. Drews, Dag Ahren, O. Hellgren

## Abstract

The malaria parasite *Plasmodium relictum* is one of the most widespread species of avian malaria. As is the case in its human counterparts, bird *Plasmodium* undergoes a complex life cycle infecting two hosts: the arthropod vector and the vertebrate host. In this study, we examine the transcriptome of *P. relictum* (SGS1) during crucial timepoints within its natural vector, *Culex pipiens quinquefasciatus*. Differential gene-expression analyses identified genes linked to the parasites life-stages at: i) a few minutes after the blood meal is ingested, ii) during peak oocyst production phase, iii) during peak sporozoite phase and iv) during the late-stages of the infection. A large amount of genes coding for functions linked to host-immune invasion and multifunctional genes was active throughout the infection cycle. One gene associated with a conserved *Plasmodium* membrane protein with unknown function was upregulated throughout the parasite development in the vector, suggesting an important role in the successful completion of the sporogonic cycle. Transcript annotation further revealed novel genes, which were significantly differentially expressed during the infection in the vector as well as upregulation of reticulocyte-binding proteins, which raises the possibility of the multifunctionality of these RBPs. We establish the existence of highly stage-specific pathways being overexpressed during the infection. This first study of gene-expression of a non-human Plasmodium species in its natural vector provides a comprehensive insight into the molecular mechanisms of the common avian malaria parasite *P. relictum* and provides essential information on the evolutionary diversity in gene regulation of the Plasmodium’s vector stages.

## Introduction

*Plasmodium* parasites are best known due to the dramatic mortality and morbidity they cause in humans across the Southern haemisphere. This group of parasites can also be found infecting a diverse range of other hosts including non-human primates, bats, rodent, reptiles, and birds (Ott 1967; Levine 1988). But for a few minor differences(Schall 1996), all these species share a nearly identical life cycle, with an asexual replicative stage in the vertebrate host, and an sexual stage a blood-sucking culicid mosquito (Diptera: Culicidae). The avian *Plasmodium* clade includes some of the world’s most genetically diverse (Bensch, et al. 2009) and virulent (Warner 1968) of all malaria parasites known to date (Valkiūnas 2005; Chagas, et al. 2017) and shows a large differentiation both in its geographical and host distribution (Medeiros, et al. 2013; Chagas, et al. 2017).

Avian malaria parasites have played a key role in our comprehension of the prevalence, morbidity and epidemiology of the disease in natural populations (Sylvia M. Fallon, et al. 2005; Valkiūnas 2005). Due to their high prevalences, widespread distributions and diverse host ranges, they have also been used to address several evolutionary issues such as host-parasite co-evolution (Charleston and Perkins 2003; Mu, et al. 2005), virulence evolution (Schall 2002; Bell, et al. 2006), and sexual selection (Spencer, et al. 2005). To date there are more than 40 morphologically described species of avian *Plasmodium* and over 1200 cytochrome b lineages, the vast majority of which are thought to be reproductively isolated entities (Bensch, et al. 2004; Bensch, et al. 2009). *Plasmodium relictum* is the most prevalent and widespread morphospecies of avian malaria and also has a highly diverse host range (Hellgren, et al. 2015), which places it amongst the top 100 most invasive species (Boudjelas, et al. 2000). This parasite species has also been found to be associated with the decline and extinction of several bird species on the islands of Hawaii (van Riper, et al. 1986; Atkinson and LaPointe 2009), with the mortality, in wild, endemic, and indigenous birds in New Zealand (Lapointe 2012) and the mortality of penguins in zoos acroos the world (Vanstreels, et al. 2015). Of the different mitochondrial cytochrome b lineages described to date within the morphologic species of *P. relictum* SGS1 the most common in terms of geographic range, host range and host prevalence. This lineage has been found to infect 129 different bird species, (MalAvi, 2020-05-08, (Bensch, et al. 2009)) and may cause severe disease and even mortality in wild birds (Palinauskas, et al. 2008; Palinauskas, et al. 2011).

All *Plasmodium* parasites share a similar life cycle requiring two infection cycles: one in the arthropod vector and one in the vertebrate host. When competent mosquitoes take an infected blood meal, they ingest both male (*microgametocytes*) and female (*macrogametocytes*) parasites. Within the first few minutes, the gametocytes transform into *gametes* and fuse within the midgut to form a diploid *zygote*. The zygotes, in turn, undergo meiosis, become motile and elongated (*ookinetes*), traverse the wall of the midgut and start to develop into *oocysts*. Over the course of several days, the oocyst undergoes several rounds of mitosis to create a syncytial cell with thousands of nuclei. In a massive cytokinesis event, thousands of *sporozoites* erupt from each oocyst and migrate through the haemocoel towards the salivary glands (Gerald, et al. 2011). Between 10-14 days after the initial infected blood meal, the parasite is in the salivary glands ready to be transmitted to a new host. When the mosquito takes a second blood meal, it injects sporozoites into the blood of the new host. These sporozoites multiply in various organs and blood cells of the host and a few days later end up releasing daughter parasites called *merozoites* into the bloodstream. Merozoites will continue the cycle of the parasite by invading other red cells and eventually produce the micro- and macrogametocytes which will restart the cycle in the mosquito (Valkiūnas 2005).

The complex life cycle of the parasites requires a considerable amount of plasticity to allow them to successfully invade a variety of widely different tissues in both of its hosts (Valkiūnas 2005; Aly, et al. 2009; Srivastava, et al. 2016). However, the degree to which this plasticity is accived through differences in generegulation of the same genesets or wheter different genes are linked to different life stages are yet to be studied. Within the mosquito, the parasite faces several developmental bottlenecks for determining the transmissibility of the parasite (Sinden, et al. 2007; Vaughan 2007; Aly, et al. 2009; Akinosoglou, et al. 2015): 1) the transition between the ingested micro (male) and macro (female) gametes to the formation of motile ookinetes, 2) the transition between the ookinete to the the oocysts and 3) the transition between oocysts to the sporozoites in the salivary glands (Valkiūnas 2005). The molecular mechanisms underlying the different developmental stages of the parasite within the mosquito have to date only been studied in human malaria (Lindner, et al. 2019) and a few species of rodent malaria species using a non-natural mosquito vector (Xu, et al. 2005). Therefore, in order to be able to find and study traits and genetic mechanisms that either have been conserved throughout the genera of *Plasmodium* or finding uniqe mechanisms linked to host or vector specificity, there is a need for knowledge of the parasite’s molecular mechanisms in the mosquito ouside the limited group of parasites that have been studied to-date (Bozdech, et al. 2003; Otto, et al. 2010; Siegel, et al. 2014; Akinosoglou, et al. 2015; Videvall, et al. 2015; Srivastava, et al. 2016; Videvall, et al. 2017)

Here, we set out to identify and study the first full transcriptome profile of a non-mammalia malaria *Plasmodium* as it goes trough its different life stages in its natural mosquito vector. First we carry out an experiment to establish the temporal dynamics of *Plasmodium relictum* (SGS1) development within its natural vector, the mosquito *Culex pipiens quinquefasciatus*. With this data in hand, we then used RNA sequencing to obtain the parasite’s transcriptome profiles at key points of its development within the mosquito, namely: 1) in the first minutes after ingestion (30 minutes post infection on average, henceforth 30mpi), 2) during the peak oocyst production phase (8 days post infection, henceforth 8dpi), 3) during the peak sporozoite production phase (12dpi) and 4) during the latest stages of the mosquito infection when the sporozoites are mainly present in the salivary glands(22dpi). As a reference point, we also obtain the transcriptome of an infected blood sample from the host bird taken immediately before the mosquito feed. We discuss how our observations will open up avenues of investigation into the pathways and molecular mechanisms that have been evolutionary conserved since the mammalian-avain malaria split.

## Results

### Temporal dynamics of *Plasmodium* development in mosquitoes

The temporal dynamics of *Plasmodium* inside the mosquito were consistent between mosquito groups fed on the three infected birds (Figure S1). Oocysts first become visible in the mosquito midgut on days 4-6 after the blood meal and the peak oocyst burdens were reached on days 8-10 (Figure 1, Figure S1). Thereafter a rapid decrease in the number of oocysts was observed so that by day 12 post blood meal, only between 2-20% of the peak oocyst burden remains. The first sporozoites appear on the head-thorax homogenate on days 8-10, only 2-4 days after the first oocysts. Sporozoite concentrations reach the peak on day 12 (Figure 1), except for the mosquitoes fed on the bird with the lowest parasite load (Bird_3, Figure S1) where the peak is reached much later (day 16, Figure S1). Onwards the number of sporozoites decreases steadily but are still detectable, albeit at low numbers, three weeks after the initial infection (Figure 1).

**Figure 1:**
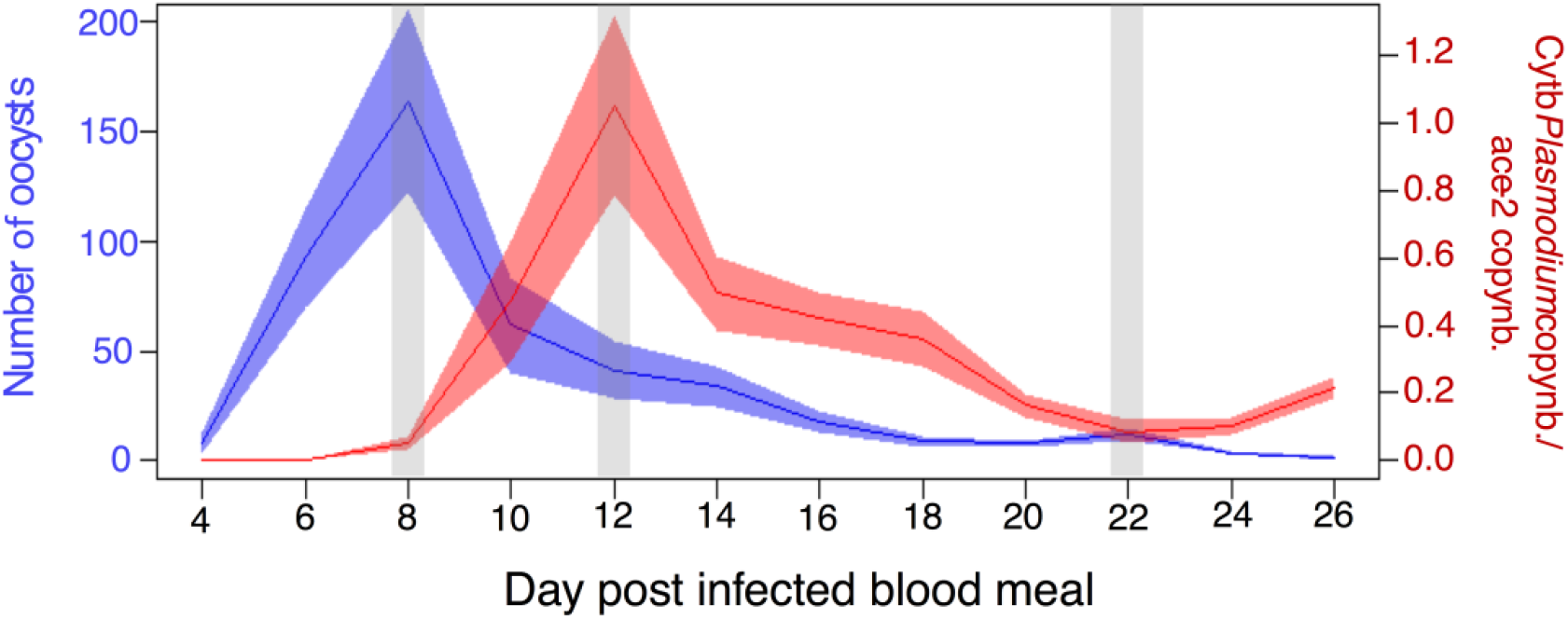
Temporal dynamics of *Plasmodium* development in mosquitoes. Average oocyst burden (blue) and sporozoite (red) counts at each dissection day. Blue and red shadows represent standard error. The left axis represents the average number of oocysts counted per female. The right axis represents the amount of sporozoites quantified by qPCR as the ratio of the parasite’s *cytb* gene relative to the mosquito’s *ace-2* gene. Light grey areas represent three of the five sampling points used to study the *Plasmodium* transcriptome.

### Data preprocessing and Sequence Alignment

We assessed the sequence data for quality and contamination. FastQC did not report any sequences having an overall Phred Score lower than 24, however, six samples had a drop in the quality for last 8-18 bases (Figure S2), which were trimmed using Trimmomatic. FastQ Screen mapped the reads with an average of 13% to the Zebra finch, 20% to Canary and 91% to the malaria vector (mosquito) reference sequences. Majority of the reads from the blood samples obtained from infected bird immediately before the mosquito feed mapped to the bird reference sequences (**Figure S3A**; Table S2). After removal of reads that uniquely mapped to the birds and vector genomes, an average of 2% of the remaining reads mapped to the parasite genome (around 0.65Million reads) and were retained for further analysis (Figure S3B;Table S3). On average 71% of paired-end reads were retained after trimming for low-quality reads and adapter content (Table S4). GC content-based filtering for the transcriptomic data was not performed as it has been estimated that the GC percentage of coding sequences (CDSs) are greater than the genomic GC percentage in the *Plasmodium* spp. (Yadav and Swati 2012). The average GC content and average length of sequences processed by FastQ Screen and Trimmomatic were 37% and 138bp, respectively (Table S5). HISAT2 produced an average of 77% overall alignment for the transcriptome data aligned to *P. relictum* genome (Table S6).

### Transcript assembly and Differential Gene Expression Analysis

Transcript assembly and abundance calculation reported 5286 genes (that we found an higher amount of genes than present is due to that alternative splice variants is interpreted as different genes) in the gene count table, which include mitochondrial and apicoplast genes (Table S7). Of these, 160 of the genes found in the annotated genome had no expression data in any of the samples and were dropped from further analysis as they did not contribute to any information to the analysis. The remaining 5126 genes were analyzed for differential gene expression using the DESeq2 package in the R statistical suite. A principal component analysis of expression clustered parasite samples belonging to the same time point together (Figure 2A). The samples are separated on PC1 (53% variance), which explained the time from the start of infection and is indicative of parasite growth (30mpi -> 8dpi -> 12dpi -> 22dpi). The samples clustered around the given timepoints indicating that the biological replicates displayed a similar expression profile.

**Figure 2A:**
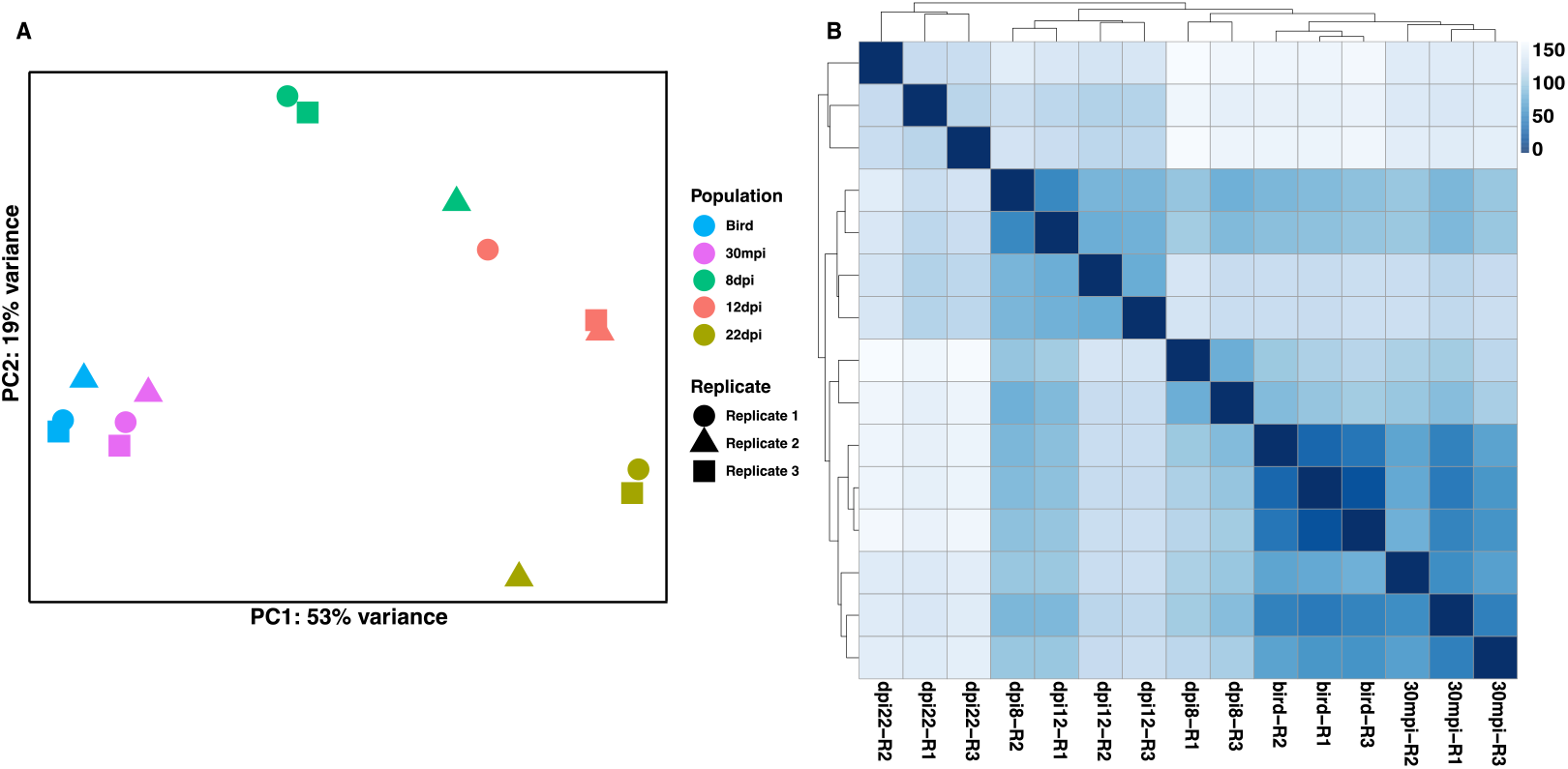
PCA plot with variable stabilizing transformation. The samples cluster together around each the time point (different stages of parasite life cycle in the mosquito). However, one 8dpi sample (from replicate 2) and one 12dpi sample (from replicate 1) cluster close to each other and away from their corresponding time point. These results are supported supported by the heatmap (B). **2B: Heatmap portraying Euclidian distance measured between the different samples.** Lighter color indicates greater distance.

Four time points (30mpi, 8dpi, 12dpi, and 22dpi) were compared for *P. relictum* genes differentially expressed against the bird samples. As baseline control, we used the transcriptomic profile of the parasite in the vertebrate host immediately before the blood meal (bird samples). The analysis reported 71, 311, 605 and 421 significantly differentially expressed genes for each of the time points (Table 2, Table S8-S11). Over 60% of these genes were found to be downregulated (Figure 3; Figure S6). Genes that were significantly downregulated with respect to the baseline, were interpreted as being highly expressed genes that are linked to the development within the vertebrate host. (Figure 4). For this study, we therefore only considered the genes that were significantly upregulated within the mosquito as compared to our baseline controls.

**Figure 3:**
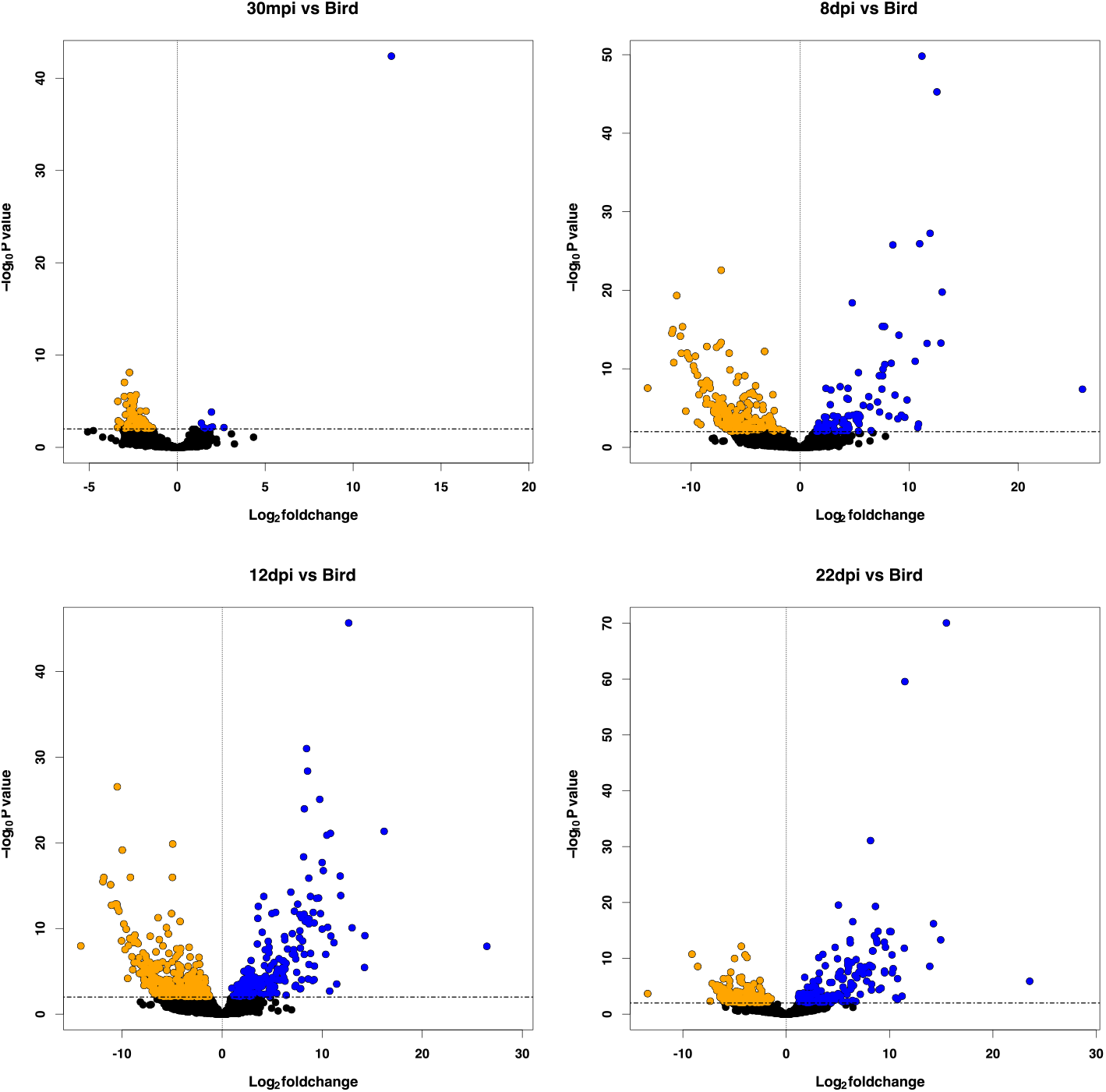
Volcano plots showing log2 fold change in expression on the x-axis and P-adjusted values on the y-axis for each of the 4 time points. Each dot represents a different gene. Differentially expressed genes with an adjusted p-value < 0.01 and absolute fold change > 0 are colored blue (upregulated) and differentially expressed genes with an adjusted p-value < 0.01 and absolute fold change < 0 are colored orange (downregulated).

**Figure 4:**
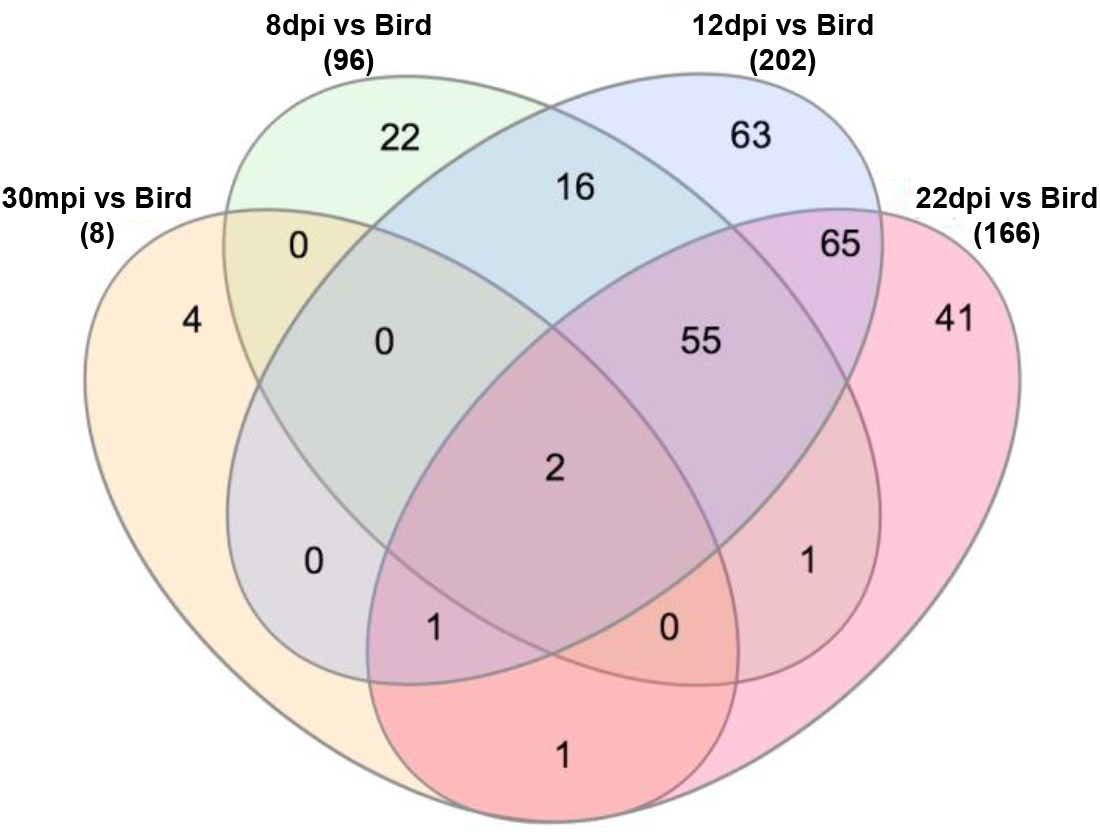
Venn diagram representing a crosswise comparison of upregulated genes in each of the 4 time points. Orange (30mpi), Green (8dpi), Blue (12dpi) and Red (22dpi). 2 genes are upregulated during all stages of infection versus Bird whereas 4, 22, 63 and 41 genes are exclusively upregulated during the 30mpi, 8dpi, 12dpi and 22dpi timepoints versus Bird, respectively.

**Table 1.** Experimental setup and sample details

| Sample ID | Sample source | Bird number | Timepoint |
| --- | --- | --- | --- |
| 1 | Mosquito | Bird 1 | 8 days post infection |
| 2 | Mosquito | Bird 1 | 12 days post infection |
| 3 | Mosquito | Bird 1 | 22 days post infection |
| 4 | Mosquito | Bird 2 | 8 days post infection |
| 5 | Mosquito | Bird 2 | 12 days post infection |
| 6 | Mosquito | Bird 2 | 22 days post infection |
| 7 | Mosquito | Bird 3 | 8 days post infection |
| 8 | Mosquito | Bird 3 | 12 days post infection |
| 9 | Mosquito | Bird 3 | 22 days post infection |
| 10 | mosquito+blood* | Bird 1 | 30mins post infection |
| 11 | mosquito+blood* | Bird 3 | 30mins post infection |
| 12 | mosquito+blood* | Bird 2 | 30mins post infection |
| 13 | Blood | Bird 1 | Infected bird |
| 14 | Blood | Bird 2 | Infected bird |
| 15 | Blood | Bird 3 | Infected bird |

\*sample contain un-digestated bird blood

**Table 2.** Number of Differentially Expressed Genes (DEG) in each case

| Case | All DEG | DEG upregulated | %Upregulated DEG |
| --- | --- | --- | --- |
| 30mpi vs Bird | 71 | 8 | 11.2 |
| 8dpi vs Bird | 311 | 96 | 30.8 |
| 12dpi vs Bird | 605 | 202 | 33.3 |
| 22dpi vs Bird | 421 | 166 | 39.4 |

Two genes were reported to be upregulated at all time points versus Bird. One of these genes had a known function associated with 28S ribosomal RNA and the other one was a conserved *Plasmodium* membrane protein with unknown function. The genes involved in sporozoite invasion, TRAP-like protein, early transcribed membrane protein, cysteine repeat modular protein, oocyst capsule protein Cas380, p25-alpha family protein, Circumsporozoite protein (CSP) were upregulated in all stages except the 30mpi stage. Two reticulocyte binding proteins were upregulated during all stages except the 30mpi stage. Several other conserved *Plasmodium* proteins with unknown function were also reported to be upregulated across timepoints. These results are summarized in Table 3, and detailed information about the corresponding genes is included in the Supplementary File (Table S12).

**Table 3.**
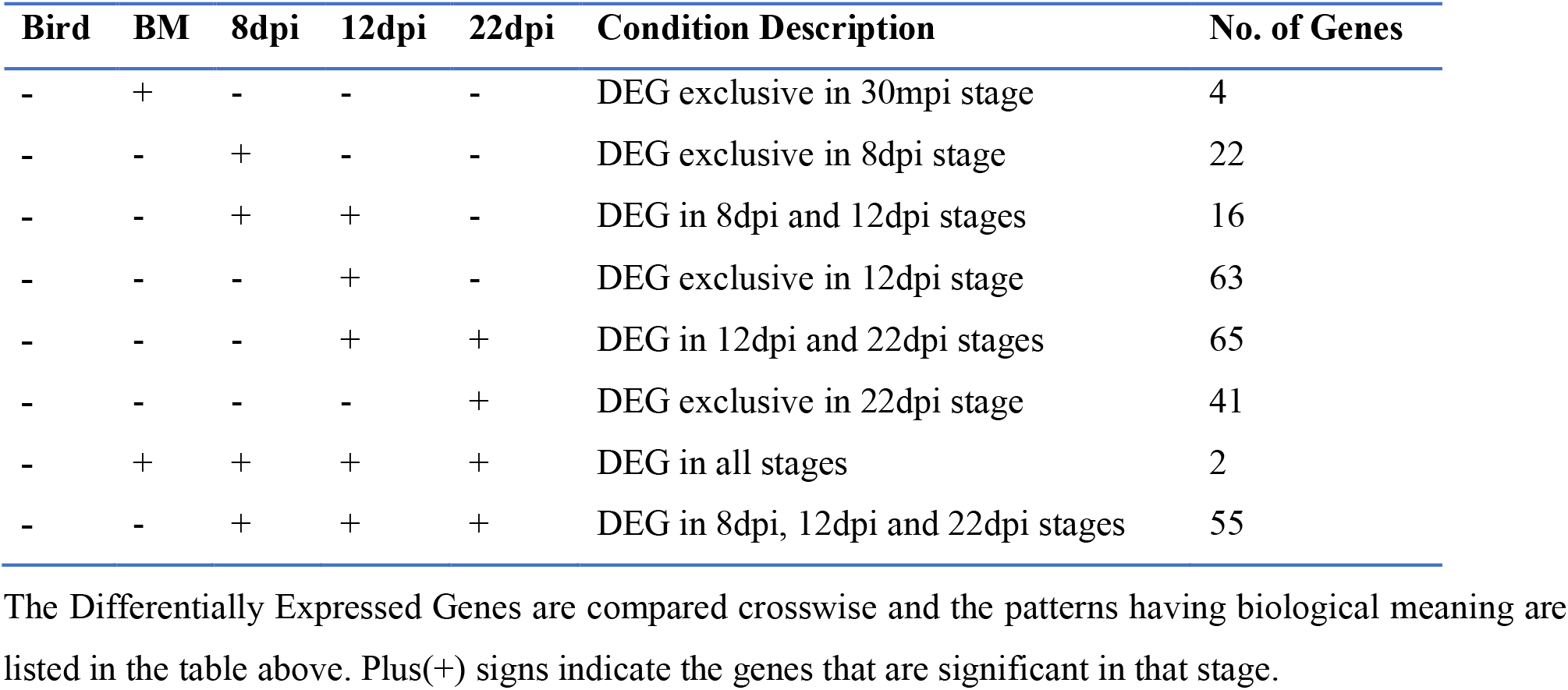
Number of Differentially Expressed Upregulated Genes (DEG) specific to each condition

### Gene Ontology Enrichment

Annotations for 2583 *P. relictum* genes were retrieved from PlasmoDB database. Gene ontology (GO) enrichment analysis revealed distinct GO enriched at different time points (Figure 5; Figure S7; Figure S8). GO terms associated with pseudouridine synthesis were enriched for the 30mpi time point, several GO terms related to oxidoreductase activity were enriched for 8dpi, GO terms related to rhoptry neck, entry into host cell (vector in our study) and membrane part were enriched for 12dpi, and GO terms related to locomotion, regulation of RNA metabolic process and lipid metabolic processes were enriched for 22dpi. GO terms related to DNA-binding transcription factor was enriched from 8dpi to 22dpi. GO terms related to exit from host, movement in host environment (vector in our study) and protein kinase activity were enriched during the late stages of infection (12dpi and 22dpi). Most of the significantly overexpressed GO terms were supported by only a few genes (Table S13-S16).

**Figure 5:**
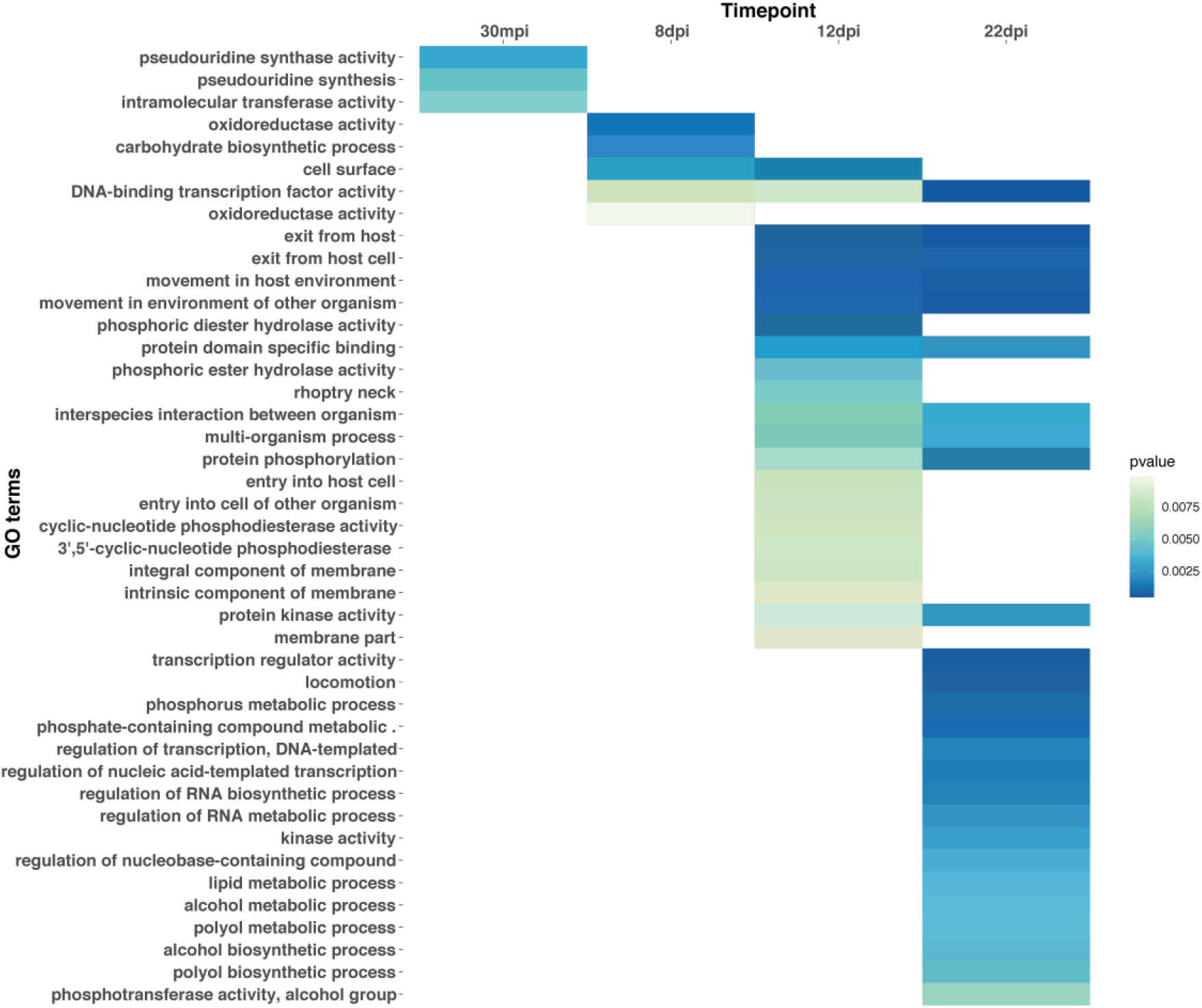
Gene ontology enrichment analysis for upregulated DEGs. Simplified figure shows the GO terms significantly enriched (pvalue <0.01) for different biological processes, molecular functions and cellular components for each time point as compared to the bird baseline. Dark color indicates higher pvalue.

## Discussion

*Plasmodium* undergoes complex molecular processes during its lifecycle to successfully invade the vector and overcome its defence mechanisms to continue the infection cycle. However, our knowledge about the processes, pathways and enzymes involved during these stages in non-human malaria parasites is limited to studies of rodent malaria parasites using a non-natural vector (Xu, et al. 2005). Given the well-documented differences in parasite performance between natural and non-natural *Plasmodium*-mosquito combinations (Aguilar, et al. 2005; Cohuet, et al. 2006; Jaramillo-Gutierrez, et al. 2009), these latter results have to be interpreted with caution. In this study, we study the transcriptome of *Plasmodium relictum* in its natural vector: the *Culex pipiens* mosquito. The average transcriptome GC content for the reads retained after quality control and that mapping to the *P. relictum* genome was calculated to be higher than that of *P. falciparum* and the avian malaria genome of *P. ashfordi* (Videvall, et al. 2017) (Figure S4). This could be due to the variation in codon usage bias and gene regulation mechanisms between the species as explained by Yadav *et al*. (Yadav and Swati 2012). Here, we computed the correlation between the duplication level and GC content (Figure S5) but did not investigate this issue any further. The PC analysis (Figure 2A) clustered the samples belonging to the same time point together, with a clear separation along the PC1 axis, as expected when same time points share similar expression profiles. However, two samples, one from 8dpi and one from 12dpi, corresponding to the peak oocyst and peak sporozoite formation (Figure 1), clustered half-way between these two time points, indicating an active transition stage. The fact that these two samples originate from the same individual bird, suggests that this is due either to a delay (8dpi) or an early onset (12dpi) of the infection in the mosquitoes due to differences between the birds. These differences could stem from differences in bird parasitaemia which are known to influence gene expression of the parasites in the host (Videvall, et al. 2017). We note that both of these intermediate samples come from the bird with the highest parasitaemia (bird #2: 7.33%, birds #1 and #3 had 5.12% and 4.48%, respectively).

### 1. Invasion genes are expressed throughout the sporogonic cycle

The majority of the genes in the parasite genome, apart from 160 genes, are expressed at one stage or other in the parasite’s life cycle within the mosquito. The differential gene expression analysis revealed that less than 40% of the total genes were upregulated at a given time point during the infection cycle in the vector as compared to our baseline control. The transition from gametocyte activation in the mosquito midgut (30mpi) to the salivary gland invasion by sporozoites (8dpi to 22dpi, Figure 1) is associated with the expression of several important genes specific to mobility, entry into the host, DNA transcription and pathogenesis (Table S12). These include a putative sporozoite invasion associated protein, which is reported to participate in host-pathogen interactions during cell-traversal (Arévalo-Pinzón, et al. 2011), a Thrombospondin-related protein 1 and a Cysteine repeat modular protein 4 (CRMP4) which facilitate the salivary gland invasion. These genes have been found in previous human malaria (Wengelnik, et al. 1999; Thompson, et al. 2007) and rodant malaria studies (Sultan, et al. 1997; Thompson, et al. 2007; Douradinha, et al. 2011) indicating that these genes are evolutionary conserved. The oocyst growth and the sporozoites release are concomitant during a long period of time (Figure 1). We observe several genes which are associated with sporozoite development to be upregulated throughout the 8dpi-22dpi transition. Consistent with previous reports (Warburg, et al. 1992; Ménard, et al. 1997b; Wang, et al. 2005; Coppi, et al. 2011; Aldrich, et al. 2012), our analysis revealed Circumsporozoite protein (CSP) to be upregulated during these stages. CSP is a key gene having a multifunctional role in oocyst development, formation of sporoblast and sporozoites, salivary gland infection, onset of sporozoites invasion, sporozoite mobility, salivary gland invasion and hepatocyte infection (Warburg, et al. 1992; Ménard, et al. 1997a; Wang, et al. 2005; Aly, et al. 2009). Interestingly, three genes associated TRAP-like proteins (TLP) were also found to be upregulated during these stages, which is in line with studies that have reported TLP to an important player in sporozoite cell traversal (Moreira, et al. 2008). Oocyst capsule protein Cap380, reported to be active during the oocyst development, sporozoite differentiation and distruption of this genes may affect parasite’s ability to invade host or vector cells (Srinivasan, et al. 2008), was also in our study upregulated during this transition highlighting its essential role in the parasite growth within the vector. In contrast to previously reported, two Reticulocyte-Binding Proteins (RBPs) which are involved in red blood cell invasion (Videvall, et al. 2017), were upregulated during 8dpi-22dpi transition. Due to sequence divergence, we cannot know with certainty which of the RBPs is the orthologous gene to the RBP we identified in this study. This suggests, either that these genes, which play a crucial role in the blood stages of the parasite, have pleiotropic effects in the mosquito, or that they are RBPs-like genes that have acquired another function in the avian malaria system.

### 2. Gene expression during the different stages in vector

Along with the number of invasive genes, several genes were specifically upregulated during a certain timepoint (Table S12).

#### 2.1. During 8dpi and 12dpi

Secreted ookinete protein 25 was upregulated at both these time point (8dpi and 12dpi) which are characterised by peak oocyst formation and peak sporozoite concentration, respectively. Secreted ookinete protein 25 is reported to affect ookinete formation and the formation of midgut oocysts (Zheng, et al. 2017). Other genes that are upregulated during this transition are genes involved in motility, *Plasmodium* exported protein (PHIST), asparagine-rich antigen, neurotransmitter, oxidation-reduction process and surface related antigen SRA.

#### 2.2. During 12dpi and 22dpi

Genes involved in sporozoite maturation and cell invasion, copies of erythrocyte membrane-associated antigen and reticulocyte binding proteins were also upregulated during the later stages of infection (12dpi to 22dpi). These include sporozoite surface protein 3, Thrombospondin-related anonymous protein (TRAP), CRMP1, CRMP2 and Calcium dependent protein kinase 6 (CDPK6). RBPs are essentially involved in erythrocyte invasion (Videvall, et al. 2017) and along with Sporozoite surface protein, CRMP1 and CDPK6, effectively invade the salivary gland during the later stages of the infection (Mota, et al. 2002; Aly, et al. 2009). Sporozoite micronemal protein essential for cell traversal (SPECT1), a key gene involved in host cell traversal (Mota, et al. 2001), are also upregulated during this transition. TRAP is a key player in the attachment and parasite invasion of the salivary glands (Sultan, et al. 1997). Other interesting genes upregulated during this transition are putative plasmepsin X, which is involved in entry and exit into the host cell, and several copies of serine/threonine protein kinase and eukaryotic translation initiation factor 2-alpha kinase 2, which are involved in ATP binding and protein kinase activity and 6-cysteine protein which affects the sporozoites’ ability to invade host cells.

#### 2.3. During 30mpi

Our analysis also reported genes which are exclusive to each time point (Table S12). We identified 4 genes upregulated exclusively during the 30mpi stage. One such gene, Putative schizont egress antigen-,1 is exclusively upregulated and is amongst genes upregulated only at this time point. Studies with *Plasmodium falciparum* have reported that inhibiting this gene prevents the schizonts from leaving the infect RBCs thereby affecting infection (Raj, et al. 2014). Whether this affects the zygote production by reducing differentiation of male and female gametocytes is an interesting question to explore. Three other conserved *Plasmodium* protein with unknown functions were also upregulated during this stage suggesting a direct or indirect role in successful parasite infection and/or zygote development.

#### 2.4. During 8dpi

At the 8dpi time point, we observed an upregulation of several genes involved in translation initiation: ATP dependent RNA helicase DBP1, and eukaryotic translation initiation factor 4E. We also found several genes associated with carbon and TCA cycle. These included NADP-specific glutamate dehydrogenase, succinyl-CoA ligase, phosphoenolpyruvate carboxylkinase. Previous studies in *P. falciparum* have reported that carbon and TCA cycle are critical for oocyst production and maturation (Srivastava, et al. 2016). We also identified a putative gene, coproporphyrinogen-III oxidase know to be associated with oxidation-reduction processes which are known to be critical processes for parasite growth under stress condition such as against the host defence mechanism in mosquito vector. As observed in the temporal dynamics of the parasite development in the vector, our analysis supports the hypothesis that this stage is marked by the peak oocyst production in the infection cycle and suggests that these genes might have a critical role in cell regulation metabolism in *Plasmodium* species.

#### 2.5. During 12dpi

Several genes involved in oocyst rupture were reported to be significantly upregulated exclusively during the stage which is marked by the peak of oocysts burst and sporozoite release (12dpi). These included oocyst rupture protein 1 and oocyst rupture protein 2 (Siden-Kiamos, et al. 2018). Genes involved in host cell invasion, such as putative rhoptry neck protein (RON6), merozoite surface protein 1 paralog and rhomboid protease ROM1 were also found upregulated during this stage. Other upregulated genes included lysine-specific histone demethylase, which is associated with oxidoreductase activity, putative inorganic anion exchange, involved in sulfate transmembrane transporter activity, and cGMP-specific phosphodiesterase. A previously reported protein involved in mobility and infectivity of the ookinetes, CDPK3 (Siden-Kiamos, et al. 2006), and one conserved *Plasmodium* protein associated with microtubule based movement was also upregulated along with several other conserved *Plasmodium* proteins with unknown functions, suggesting their role in oocyst maturation and rupture, sporozoite formation, release and transportation.

#### 2.6. During 22dpi

The last time point (22dpi) characterised by the low density of sporozoites. During this time, the sporoozites are mainly present in the salivary glands. Putative sporozoite and liver stage asparagine-rich protein (SLARP) and CDPK5 are upregulated during this timepoint. These proteins which are reported to have a role in the regulation of transcription, have been previously described in *Plasmodium falciparum* (Silvie, et al. 2008; Aly, et al. 2011). Upregulation of these genes and other genes involved in kinase activity – serine/threonine protein kinase, inositol polyphosphate multikinase, suggests an essential role in invasion mechanism during this transition. Some of the previously reported stage specific markers identified in other *Plasmodium* species was not observed in our study, such as UOS3 (Mikolajczak, et al. 2008; Combe, et al. 2009; Steinbuechel and Matuschewski 2009), MAEBL (Kariu, et al. 2002), CelTOS (Kariu, et al. 2006). If this pattern is due to methodolocical reasons or a true biological difference we cant determine at this point but is important to disentangle in future studies in order to understand how the gene functions have evolved across the different species.

### 3. Role of AP2 transcription factors in the sporogonic cycle

Several copies of putative AP2 domain transcription factors are active during the infection, and our analysis also captures a few specific AP2 transcription factors associated with the various stages of parasite development. Previous studies have reported the functional role of AP2-SP2 during the sporozoite formation (Yuda, et al. 2010). In our analysis, we identified this transcription factor to be upregulated from 8dpi to 22dpi. Previous finding report that AP2-O3 is associated with ookinete formation, gliding and invasion (Modrzynska, et al. 2017). However the direct target of this transcription factor is unknow. We observe AP2-O3 (putative) to be upregulated at 8dpi, which might suggest that AP2-O3 regulates genes during the peak oocyst stage. Putative AP2-EXP which, in *P. falciparum*, AP2-EXP is reported to regulate virulence genes (Martins, et al. 2017), is upregulated exclusively at 12dpi. Our results are consistent with previous studies in *P. berghei* (Yuda, et al. 2010) where AP2-EXP has been shown to be expressed specifically during the sporozoite stage. AP2-L, an AP2 transcription factor associated with liver exoerythrocytic stages (Iwanaga, et al. 2012) and sporozoites (Yuda, et al. 2010) was also found to be upregulated during the 22dpi stage of infection. We also identified upregulation of AP2-O5 during this stage. Even though the role of AP2-O in activation of genes associated with ookinete mobility and oocyst development has been previously established (Yuda, et al. 2009), the exact function of AP2-O5 during the sporozoite stage is unclear. Furthermore, several copies of putative AP2 transcription factors were upregulated throughout the duration of infection suggesting that these transcription factors might be responsible for regulation of specific genes in the respective stages of infection. This could be explained by the fact the experiment is conducted *in vivo* and that the different parasite development stages are not completely separate in time. Many copies of AP2 genes indicate that there are several transcriptional factors that are associated and active at different time points, indicating the strong importance of these transcription factors during the entire infection process. This also suggests that AP2 genes might have very selective roles in the parasite development processes and are tightly regulated. The presence of these essential genes in distantantly related mammalian and avian *Plasmodium* also suggest that they share some essentially conserved genes (Aly, et al. 2009).

### 4. GO enrichment revealed highly stages specific pathways and metabolism

The different *Plasmodium* species have diverged over time and several genes might have been lost or aquired during the evolution. Many of the genes in the *P. relictum* genome are uncharacterized (Böhme, et al. 2018). This potentially limits our ability to pin down all the essential mechanism during the lifecycle of the parasite. To strengthen our understanding of the biological functions behind some of the transcripted genes, we make use of Gene Ontology (GO) system of classification for the genes. GO enrichment allows to establish if the genes of interest are associated with certain biologicalbased on statistical testing. For this purpose, we used the up-to-date annotation for *Plasmodium relictum* from PlasmoDB which includes both experimentally validated and computationally predicted GO terms. We observed several GO terms enriched in a very time specific manner suggesting the coordinated regulation of these processes during the development of the parasite (Table S13-S16).

#### 4.1. 30mpi

Only a few GO terms were significantly enriched for the upregulated genes in 30mpi stage. These included pseudouridine synthesis, pseudouridine synthesis activity and intermolecular transferase activity. This is indicative of an active progress during this stage of gamete maturation and zygote formation. The successful production of zygotes and the ookinete formation is supported by the overexpressed pseudouridine synthesis which is directly associated with macromolecule modification. Surprisingly, we also saw the GO term associated with reproduction (GO:0000003) being reported amongst the top 20 GO terms enriched for biological processes. This term was, however, statistically not significant in our analysis (Figure 5; Table S13), probably due to the limitation in the available knowledge regarding the genes annotated to this term.

#### 4.2. 8dpi

At 8dpi the repeated nuclear division (endomitosis) in oocysts is almost at its peak. This stage is enriched for GO terms associated with oxidoreductase activity, carbohydrate biosynthesis process, DNA binding transcription factor and cell surface (Figure 5; Table S14). Consistent with DGE analysis, transcription and metabolic processes are active and essential for energy demand during parasite growth and transition (Srivastava, et al. 2016). This analysis also reveals the overexpression of oxidation-reduction process which supports regulation of redox balance in the parasite. During infection, the parasites can fight the host’s immune oxidative stress mechanism by altering the redox balance and antioxidant defence system (Becker, et al. 2004; Müller 2004).

#### 4.3. 12dpi

The 12dpi stage is associated with overexpressed GO terms associated with the exit from the host cell, movement in the host environment, interspecies interactions, multi-organism processes (meaning interaction between organisms), protein phosphorylation, and entry into the host cells. The overexpression of these GO terms along with overexpression of GO terms associated with cell surface, membrane parts and integral component of membrane, reflects the crucial role of these pathways in toocyst maturation, sporozoite release and invasion (Patzewitz, et al. 2013)(Figure 5; Table S15). Other GO terms relating to the interaction with the host, the formation of the parasite’s cellular and its mobility within the host seem to be critical for the parasite entry and transmission (Aly, et al. 2009). Enrichment of the GO term rhoptry neck indicates a host cell invasion mechanism active during this transition. Several molecular functions related to protein domain specific binding, DNA binding transcription factor activities and a number of protein kinase activities are also enriched during this stage, which are indicative of active transition from oocyst to sporozoite formation.

#### 4.4. 22 dpi

A wide range of biological processes are associated to the genes upregulated during the later stage of the infection (22dpi). The overexpressed GO terms are associated with the exit from host cell (vector cells in our study), movement in the host environment (vector cells in our study), protein phosphorylation, regulation of RNA metabolic processes, interaction between organisms, regulation of transcription, interspecies interaction and regulation of RNA metabolic process. The molecular functions that are overexpressed are associated with DNA-binding transcription factors, and transcription regulator, protein kinase and phosphotransferase activities (Figure 5; Table S16). Previous studies have shown the importance of parasite protein kinases throughout the growth and development of the parasite, reviewd in (Doerig, et al. 2008), and our analysis also suggests that these pathways have an essential role in gene regulation for sporozoite invasion. It is suggested that these pathways have a key role in the stage specific development of the parasite. However, several of the interesting GO terms were not significant according to our cutoff (p<0.01)(Figure S8). Our analysis were also heavily influenced by the existing GO annotation of the genes. Hence these observations have to be considered with caution. Since not a lot is known about the metabolism and enzymes in these stages, this work provides first-hand insight into the mechanism of the parasite in the vector system.

### Conclusion

This is the first study to provide a comprehensive insight into the molecular mechanism of one of the most harmful avian malaria parasite *P. relictum* in *Culex quinquefasciatus*, its natural vector thereby providing valuable knowledge about the genes involved in critical transitions in the lifecycle of the parasite. We have captured a snapshot of genes associated with host immunity, even the one lowly expressed, during at all stages during infection within the vector. We identified several known genes associated with cell invasion along with several gene with unknown functions specific to different infection stages of the parasite life cycle, which can be potential candidates for functional studies. We also identified copies of reticulocyte binding proteins upregulated during different stages of the infection, either suggesting a that these proteins have a wider function than previously thought or that they have evolved a different function in *P. relictum*. We also identified many significant genes specific to different stages of parasite development in this analysis. Although only upregulated genes were considered for this study; we acknowledge that downregulated genes could have a potential cascading effect on the parasite growth which may have been missed in our analyses. Some of the genes have been studied using genetic knockout mutants and their functions have been validated experimentally. However, a large portion of the parasite genome remains uncharacterized *Plasmodium* genes, which limits our ability to list all active genes during the sporogonic cycle. Future research will help determine the function and features of these genes. This work also contributes to improve the genetic resources for *P. relictum*. The knowledge gathered from this study could contributes to our understanding of the critical stages in *Plasmodium* life cycle and could be used as an active model to further conduct specific studies on targeted genes related to invasion. As a result, this work provides us with an insight into the functional relatedness and mechanism of the development of different *Plasmodium* species within the vector. This can inform further research aimed at devising broader ecological solutions regarding the disease control and conservation programs.

## Methods

### Temporal dynamics of *Plasmodium* sporogony protocol

To establish the temporal dynamics of *Plasmodium relictum* (SGS1) development within its natural vector, the mosquito *Culex pipiens quinquefasciatus*, experiments were carried out in three birds (*Serinus canaria*) which were infected using standard infection protocols (Pigeault, et al. 2015). Ten days later, at the peak of the acute infection stage, each bird was placed overnight in a separate experimental cage with 180 7-day old female mosquitoes (Vézilier, et al. 2010). After the mosquitoes had taken a blood meal, the birds were removed from the cages and the mosquitoes were supplied with 10% *ad libitum* sugar solution until the end of the experiment. Every two days, starting on day 4 and finishing on day 26 post-blood meal, twelve mosquitoes were haphazardly sampled from each cage. Each mosquito was dissected to count the number of oocysts in the midgut, and its head-thorax was preserved at −20°C for the quantification of the transmissible sporozoites. Developing oocysts were counted under the microscope (Vézilier, et al. 2010). Sporozoites were quantified using real-time quantitative PCR as the ratio of the parasite’s *cytb* gene relative to the mosquito’s *ace-2* gene (Zélé, et al. 2014).

### Parasite transcriptomic experimental protocol

In the next experiment, Mosquitoes (*Cx. pipiens quinquefasciatus*) were experimentally infected with *P. relictum* (SGS1) by allowing them to feed from 3 canaries (*Serinus canaria*) which had been previously infected using standard laboratory protocols (Pigeault, et al. 2015). The experimental protocol proceeded as follows. On the day of the experiment (day 0), immediately before the beginning of the experiment, 20μl blood from the bird’s wing was sampled. A drop of blood was used to quantify parasite load via a blood smear (as described by (Valkiūnas 2005), the rest was mixed with 500μl of Trizol (Life Technologies, Carlsbad, CA, USA) and frozen at −80°C for subsequent RNA extraction (henceforth ‘bird’ sample). Seven-day old female mosquitoes, reared using standard laboratory protocols (Vézilier, et al. 2010), were introduced into each of the cages 19 hours after the start of the experiment (150 female mosquitoes per cage). Cages were visited 30 minutes later and 20 fully-gorged resting mosquitoes were haphazardly sampled from each of the cages (henceforth ‘30 minutes post infection (30mpi)’ sample). Half of the mosquitoes were immediately homogeneised, mixed with 500μl of Trizol and frozen at −80°C for subsequent RNA extraction (1 tube per cage). The rest of the mosquitoes were individually frozen at −80°C. The bird was taken out at the same time, and the rest of the mosquitoes were left in the cage with a source of sugar solution (10%) at our standard insectary conditions (25-27°C, 70% RH). On day 8 after the start of the experiment, to coincide with the peak of oocyst production as estimated by the previous experiment, 30 further mosquitoes were randomly taken from each of the cages (henceforth ‘8 dpi’ sample). For each cage, 10 of these mosquitoes were homogeneised and conserved in 500μl RNAlater (ThermoFisher Scientific/Ambion, Waltham, USA). The procedure was repeated on day 12 to coincide with the peak sporozoite production (‘12 dpi’ sample) and on day 22 during the late stage of the infection (‘22 dpi’ sample).

### RNA extraction

RNA from the bird blood samples was extracted with a combination of Trizol LS and RNeasy Mini spin columns (Qiagen, Hilden, Germany). Homogenizing and phase separation was done according to the TRizol LS manufactures protocol resulting in an aqueous phase which was then mixed with one volume of 70% ethanol and placed in a RNeasy Mini spin column. From this point on the RNeasy Mini spin columns manufactures protocol was followed.

RNA from the 30mpi samples (collected in Trizol LS) were disrupted and homogenized using a TissueLyser (Qiagen, Hilden, Germany). Ten whole mosquitoes were collected in 500 μl of Trizol LS, the total volume of TRizol LS was adjusted to 750 μl and a 5 mm stainless steel bead was added. The TissueLyser was run for two cycles of three minutes at 30 Hz. Phase separation was done according to the TRizol LS manufactures protocol resulting in an aqueous phase which was then mixed with one volume of 70% ethanol and placed in a RNeasy Mini spin column. From this point on the RNeasy Mini spin columns manufactures protocol was followed.

RNA from the 8dpi, the 12dpi and 22dpi samples (collected in RNAlater) were extracted with RNeasy Mini spin columns following the manufactures protocol. Disruption and homogenization were done using a TissueLyser (Qiagen, Hilden, Germany). Ten whole mosquitoes were moved to a new tube and 600 μl of buffer RLT was added as well as a 5 mm stainless steel bead. The TissueLyser was run for two cycles of three minutes at 30 Hz.

The concentration of all RNA samples was measured on a Nanodrop 2000/2000c (Thermo Fisher Scientific, Wilmington, DE, USA). Dried in samples where shipped on dry ice to Novogene (Hong Kong) for mRNA sequencing. mRNA from each time point was sequenced using Illumina HiSeq platform at an average of 85 M reads per liberary. We obtained paired-end reads of 150bp length for 15 sequenced transcriptomes which were used for further analysis.

### Data preprocessing

We examined the RNA-Seq reads for quality, per-base sequence content, adapter content and overrepresented sequences using FastQC (Version 0.11.5) (Andrews, et al.). Contamination due to other genomes such as human and bird was estimated using FastQ Screen (Version 0.11.1) (Wingett and Andrews 2018). FastQ Screen maps the raw-reads to the indexed reference genome using Bowtie 2 (Langmead and Salzberg 2012) or BWA (Li and Durbin 2009) to calculate an estimate of the reference genome contamination in query sequences. We screened the transcriptome sequences against genomes of Zebra Finch (GCF_000151805.1_Taeniopygia_guttata-3.2.4) and *Serinus canaria* (GCF_000534875.1_SCA1) for bird references, mosquitoes (GCF_000209185.1_CulPip1.0) for malaria vector and *Plasmodium relictum* genome (GCA_900005765.1_PRELSG) and indexed using Bowtie2 (Version 2.3.1). Birds infected with the parasites in our study was *Serinus canaria* hence we used this genome for screening transcripts along with Zebra Finch genome which is widely used as bird model genome. The genomes were retrieved in the form of Fasta sequences from NCBI Genome browser (https://www.ncbi.nlm.nih.gov/genome). All the reads that did not map uniquely to human, bird and malaria vector reference genomes were filtered out using-filter function in FastQ Screen. Adapter sequences, ambiguous nucleotides, and low-quality sequence were trimmed off using Trimmomatic (Version 0.36) (Bolger, et al. 2014). We used paired-end mode with options -phred33 for base quality encoding, a sliding window of 4:20, the minimum sequence length of 70, trailing 3 and leading 3. We used the standard Illumina adapter sequences (option: ILLUMINACLIP Truseq3-PE.fa:2:30:10) available in Trimmomatic to screen and trim the adapter sequences from the query. The choice of parameters was made after examining the reads for quality and presence of adapter sequences. Once trimmed, the sequences were again evaluated for quality (FastQC) before proceeding to the following steps. MultiQC (Ewels, et al. 2016), which summarizes results from various tools and generate a single report, was used for illustrative purpose.

### Read alignment, transcript assembly and abundance calculation

The filtered and trimmed sequence files were aligned to *Plasmodium relictum* published genome (GCA_900005765.1_PRELSG) using HISAT2 (Version 2.1.0) (Kim, et al. 2015). All the sequence files were aligned as paired end reads and with default parameter settings. HTseq-count (Anders, et al. 2015) (parameters: -s no -t gene -i locus_tag) is used to calculate the read count for each sample and custom bash scripts were written to post-process the files to generate gene count matrix. Scripts and more detailed parameter explanation for this process can be provided upon request.

### Differential Gene Expression Analysis

The differential gene expression analysis was performed in DESeq2 (Version 1.16.1) (Love, et al. 2014) package in R. Genes with no expression across all samples i.e. genes with zero read across all samples were filtered out from the gene count table before analysing the data further. To avoid having any bias in the analysis and to examine the data further, variance stabilized transformation of counts (Anders and Huber 2010) was used to perform PCA. Plots and visualization were made using RColorBrewer, genefilter, diplyr, and ggplot2 packages in R.

Differential gene expression (DGE) was performed with the filtered gene count matrix. As DESeq2 normalizes the data within the method for library size difference, the gene count matrix was not transformed or normalized beforehand for the analysis The bird samples were considered as a reference to perform DGE analysis. The p-value for DEG was set to 0.01 and the four cases were defined for the DGE analysis. The significantly differentially expressed upregulated genes (having positive log2 fold change in our comparison) after foldchange and adjusted p-value sorting, from the four cases (30mpi vs bird, 8 days post-infection (dpi) vs bird, 12dpi vs bird, 22dpi vs bird) were considered for further enrichment analysis. The genes from all the cases were compared cross-wise to identify genes common in all stages and the genes exclusive to a certain stage of infection.

### Gene Ontology Enrichment Analysis

To look for functions that where significantly overrepresented among the upregulated genes at the different stages we conducted an Gene Ontology Enrichmnet analysis. The significantly differentially expressed upregulated genes were analysed to identify overrepresented Gene Ontology (GO) terms using the topGO package (Alexa and Rahnenfuhrer 2019) in RStudio (Version 1.0.143). TopGO allows enrichment analysis for GO terms for custom background annotation. It also allows a flexible testing framework with support to many different algorithms. The GO annotation for SGS1-like strain was downloaded from PlasmoDB (http://plasmodb.org/plasmo/) and customised to fit the input format required by topGO. The background for analysis for each case is the list of genes and their p-adjusted valued as reported by DESeq2. All the significantly differentially expressed upregulated genes from each case was analysed independently to identify the ontologies enriched specifically for each stage of the parasite’s life cycle. The enrichment analysis is performed using classical algorithm and Fisher’s exact test and the GO terms with p value < 0.01 are considered enriched. The overrepresented GO were reported from all three domains: biological processes, molecular functions and cellular components.

## Acknowledgements

This study was funded by the Swedish Research Council (grant 2016-03419) and Nilsson-Ehle foundation to O.H. A.R. was funded through the ANR-16-CE35-0001-01 (‘EVODRUG’).

## Author Contributions

O.H. planned and designed the study with input from A.R and S.G. A.R and S.G. designed and performed the mosquito experiment for the RNA sequencing. V.S. performed the bioinformatic and statistical analyses with help and advice from D.A. and O.H. R.P. performed and analyzed the experiment that determined the timing of the life stages in the mosquito. A.D. performed the molecular labwork. V.S. wrote the article with input from all authors.

## Supporting Information

**Figure S1:** Mean oocyst (blue) and sporozoite (red) counts at each dissection day for each of the three birds (A: Bird 1, B: Bird 2, C: Bird 3). Shadows represent standard error. Parasitaemias and gametocytaemias for each of the birds are as follows: Bird 1: 5.12% and 1%, Bird 2: 7.33% and 0.4%, and Bird 3: 4.48% and 0.3%, respectively.

**Figure S2:** MultiQC report from FastQC analysis before filtering low quality reads and trimming for adapters and low-quality positions. Every line represents one sample.

**Figure S3: Estimate of contamination from reference genomes in FastQ Screen.** MultiQC report from FastQ_Screen analysis. Every bar represents one sequence file. The samples in each group are in the order from left to right as follows: six samples from the bird blood transcriptome stage(bird) (paired end sequence files), next six from the blood meal stage (30mpi) (paired end sequence files), next six from 8dpi (paired end sequence files), the next six from 12dpi (paired end sequence files), and the last six from 22dpi. The light blue bar indicates uniquely mapped reads to the specific genome, the dark blue bar indicates reads multi-mapping on the same genome, and the light and dark red bars indicate read mapping to multiple genomes uniquely and at multiple sites, respectively. In the first screen (Figure S3A), the samples were mapped against Zebra finch, Canary and the malaria vector (mosquito) reference genomes. All the reads that did not map uniquely to these reference genomes were then extracted and mapped against parasite genome (Figure S3B)

**Figure S4: Per-sequence GC content after trimming and filtering.** The average observed GC% was 37% which is considerably higher than the genomic GC% of the *Plasmodium relictum* (18%).

**Figure S5:** We speculate that higher GC content is due to higher duplicate levels in the samples as indicated by spearman’s rank correlation coefficient between GC content and duplicate levels.

**Figure S6: Related to Figure 3.** MA plot showing log-transformed normalized expression values for all the genes and the shrunk-en log fold change in different cases. Each gene is represented by a dot. The genes with an adjusted p-value < 0.01 are shown in red.

**Figure S7: Related to Figure 5,** figure shows the actual GO terms significantly enriched (pvalue <0.01) for different biological processes, molecular functions and cellular components for each time point as compared to the bird baseline.

**Figure S8:** Figure shows the actual GO terms significantly enriched (pvalue <0.05) for different biological processes, molecular functions and cellular components for each time point as compared to the bird baseline.

**Table S1:** Quality statistics from FastQC before filtering for contamination

**Table S2:** Fastq_screen report from first filter reporting number of reads mapping to different genomes

**Table S3:** Fastq_screen report from the second filter reporting number of reads mapping to parasite genome

**Table S4:** Trimmomatic result after filtering for contamination reporting number of reads retained after quality and adapter trimming

**Table S5:** Quality control statistics from FastQC after trimming for low quality and adapter contents

**Table S6:** HISAT2 alignment statistics

**Table S7:** Gene count csv file generated from transcript assembly and abundance calculation

**Table S8**: Differentially expressed genes for 30mpi vs bird

**Table S9:** Differentially expressed genes for 8dpi vs bird

**Table S10:** Differentially expressed genes for 12dpi vs bird

**Table S11:** Differentially expressed genes for 22dpi vs bird

**Table S12:** Differentially expressed upregulated genes exclusive and common to different timepoints with description

**Table S13:** GO terms enriched for upregulated genes for 30mpi for biological process, cellular components and molecular functions. Terms with pvalue <0.01 are considered significant and discussed further.

**Table S14:** GO terms enriched for upregulated genes for 8dpi for biological process, cellular components and molecular functions. Terms with pvalue <0.01 are considered significant and discussed further.

**Table S15:** GO terms enriched for upregulated genes for 12dpi for biological process, cellular components and molecular functions. Terms with pvalue <0.01 are considered significant and discussed further.

**Table S16:** GO terms enriched for upregulated genes for 22dpi for biological process, cellular components and molecular functions. Terms with pvalue <0.01 are considered significant and discussed further.

## Notes

### Competing Interest Statement

The authors have declared no competing interest.

